# GSDB: a database of 3D chromosome and genome structures reconstructed from Hi-C data

**DOI:** 10.1101/692731

**Authors:** Oluwatosin Oluwadare, Max Highsmith, Jianlin Cheng

## Abstract

Advances in the study of chromosome conformation capture (3C) technologies, such as Hi-C technique - capable of capturing chromosomal interactions in a genome-wide scale - have led to the development of three-dimensional (3D) chromosome and genome structure reconstruction methods from Hi-C data. The 3D genome structure is important because it plays a role in a variety of important biological activities such as DNA replication, gene regulation, genome interaction, and gene expression. In recent years, numerous Hi-C datasets have been generated, and likewise, a number of genome structure construction algorithms have been developed. However, until now, there has been no freely available repository for 3D chromosome structures. In this work, we outline the construction of a novel Genome Structure Database (GSDB) to create a comprehensive repository that contains 3D structures for Hi-C datasets constructed by a variety of 3D structure reconstruction tools. GSDB contains over 50,000 structures constructed by 12 state-of-the-art chromosome and genome structure prediction methods for publicly used Hi-C datasets with varying resolution. The database is useful for the community to study the function of genome from a 3D perspective. GSDB is accessible at http://sysbio.rnet.missouri.edu/3dgenome/GSDB

## INTRODUCTION

The three-dimensional (3D) organization of the genome plays a significant role in many diverse biological functions and processes including gene expression [1], regulation [2,3] and transcriptional regulation [4]. Several studies of the architecture of the genome in the cell have linked genome structure to the mechanism of these functions; hence, it is essential to understand the spatial arrangement within the cell nucleus in order to fully elucidate this relation [5–7]. Early studies of the structure of the genome have relied on the use of microscopy techniques such as fluorescence in situ hybridization (FISH), a technique that employs fluorescence probes to detect the presence of a specific chromosome region and the proximity between two regions in a genome sequence [8–10]. Other microscopy methods developed to study the genome organization include stimulated emission depletion (STED) [11], stochastic optical reconstruction microscopy (STORM) [12], and photo-activated localization microscopy (PALM or FPALM) [13,14]. While these techniques have proven very useful in providing insights into the organization of the genome for DNA fragments or chromatin regions, they are limited and unsuitable for an overall view of the genome-wide inter-and intra-chromosomal relationship study of the genome within the cell nucleus [15].

In order to capture these inter- and intra-chromosomal interactions, a variety of next-generation, high-throughput sequencing technologies have emerged including: 3C [16], 4C [17], 5C [18], Hi-C [19], TCC [20] and ChIA-PET [21,22]. Out of all these techniques, the Hi-C technique has seen a particularly high usage because of its ability to comprehensively map the chromatin interactions at a genome wide scale.

A Hi-C experiment results in the generation of an interaction frequency (IF) matrix for chromosomal regions (loci) within a chromosome or between any two chromosomes in a population of cells [19,23–25]. With the advancement of the Hi-C research, sophisticated tools such as GenomeFlow [23], Juicer [26], and HiC-Pro [27] have been developed to generate IF matrices from raw sequence pair reads data [28]. Some methods represent the contact matrix in a sparse 3-column format where columns 1-2 denote the interacting loci and column 3 denotes the number of interactions (or contacts) between the corresponding loci in a Hi-C dataset [24,29,30].

Many methods have been developed for chromosome 3D structure reconstruction from chromosome conformation capture (3C) such as the Hi-C data. Generally, these data-driven methods can be grouped into three classes [31] based on how the IF is used for 3D structure construction: distance-based, contact-based and probability-based. First, distance-based methods implement the 3D structure construction through a two-step process. These methods convert the IF matrix to a distance matrix between loci based on an inverse relation observed from FISH 3D distance data [19]. An optimization function is thereafter used to infer a 3D structure from an initial random structure with the objective of satisfying the distances in the distance matrix as much as possible [24,29,32–40]. Second, contact-based methods consider each chromosomal contact as a restraint and apply an optimization algorithm to ensure that the number of contacts in the input contact matrix is satisfied in the 3D structure [30,41–43]. Third, probability-based methods define a probability measure over the IF, by constructing the structure inference problem as a maximum likelihood problem and thereafter using a sampling e.g. Markov chain Monte Carlo (MCMC) or optimization algorithm to solve the prediction problem [25,44–46]. Despite the significant progress in the methodological development in 3D chromosome and genome structure modeling and availability of a lot of Hi-C datasets, there is still no public database to store 3D chromosome and genome models for the biological community to use.

Here, we present Genome Structure Database (GSDB), a novel database that contains the chromosome/genome 3D structural models of publicly and commonly used Hi-C datasets reconstructed by twelve state-of-the-art 3D structure reconstruction algorithms at various Hi-C data resolution ranging from 25KB – 10MB. The database is organized such that users can view the structures online and download the 3D structures constructed for each dataset by all the reconstruction methods. Our database is the first of its kind to provide a repository of 3D structures and the evaluation results for 3D structures constructed from many Hi-C datasets by different Hi-C data reconstruction methods all in one place.

## MATERIALS AND METHODS

### DATASETS

Our Hi-C data is pulled from a variety of sources which we list here. Some datasets were downloaded from the Gene Expression Omnibus (GEO) database, including the Hi-C contact matrices datasets (GEO accession Number: <u>GSE63525</u>) of cell line GM12878 from Rao *et al.* [47], normalized interaction matrices for each of the four cell types - mouse ES cell, mouse cortex, human ES cell (H1), and IMR90 fibroblasts – (GEO accession Number: <u>GSE35156</u>) [48,49], and the Hi-C contact matrices datasets (GEO Accession Number: <u>GSE18199</u>) of karyotypically normal human lymphoblastic cell line (GM06990, K562)[19]. All other Hi-C datasets were obtained from the ENCODE project repository [50], and the GEO accession Number and the ENCODE ID for each dataset are available on the GSDB website. Currently, GSDB contains over 50,000 structural models of various resolutions reconstructed from 32 unique Hi-C datasets by 12 state-of-the-art 3D genome/chromosome modeling methods. More Hi-C datasets will be used to build 3D models as they are available.

### NORMALIZATION

Hi-C data normalization is an important process in 3D structure reconstruction from Hi-C data, because the raw contact count matrix obtained from 3C experiments may contain numerous systematic biases, such as GC content, length of restriction fragments, and other technical biases that could influence the 3D structure reconstruction [51–55]. Consequently, all the contact matrices were normalized prior to applying the 3D structure reconstruction algorithms. GM12878 cell line datasets were normalized using the Knight–Ruiz normalization (KR) method [53, 47], and the normalized interaction matrices downloaded from Dixon et al. [48] were normalized using Yaffe and Tanay normalization method [55]. The Vanilla Coverage (VC) technique [47] was used as the default technique to normalize all the other Hi-C datasets.

### DATABASE IMPLEMENTATION

The GSDB website interface was implemented using HTML, PHP and JavaScript, and the database was implemented in MySQL (https://www.mysql.com/). The online 3D structure visualization was done through 3Dmol viewer, a molecular visualization JavaScript library [56].

### 3D MODELING ALGORITHMS INCLUDED

We used twelve existing algorithms for the 3D structure construction. We selected a mixture of distance-based, contact-based, and probability-based algorithms [31]. We first describe the distance-based algorithms. LorDG [24] uses a nonlinear Lorentzian function as the objective function with the main objective of maximizing the satisfaction of realistic restraints rather than outliers. LorDG uses a gradient ascent algorithm to optimize the objective function. 3DMax [29] used a maximum likelihood approach to infer the 3D structures of a chromosome from Hi-C data. A log-likelihood was defined over the objective function which was maximized through a stochastic gradient ascent algorithm with per-parameter learning rate [57]. Chromosome3D [32] uses distance geometry simulated annealing (*DGSA*) to construct chromosome 3D structure by translating the distance to positions of the points representing loci. Chromosome3D adopts the Crystallography & NMR System (CNS) suite [58] which has been rigorously tested for protein structure construction for the 3D genome structure prediction from Hi-C data. HSA [34] introduced an algorithm capable of taking multiple contact matrices as input to improve performance. HSA can generate same structure irrespective of the restriction enzyme used in the Hi-C experiment. miniMDS [37] proposed an algorithm to model Hi-C data by partitioning the contact matrix first into segments and building the 3D structure bottom-up from each segment which are eventually aggregated to form a final 3D structure. ChromSDE [38] (Chromosome Semi-Definite Embedding) framed the 3D structure reconstruction problem as a semi-definite programming problem. Shrec3D [39] formulated the 3D structure reconstruction problem as a graph problem and attempts to find the shortest-path distance between two nodes on the graph. The length of a link is determined as the inverse contact frequency between its end nodes. Each fragment is regarded as the nodes connected by a link. The represented 3D structure for a Hi-C data is one in which distance between the nodes is the shortest. InfoMod3DGen [40] converts the IF to a distance matrix and used an expectation-maximization (EM) based algorithm to infer the 3D structure.

In the contact-based category, we used MOGEN [30] and GEM [42] for the 3D structure reconstruction. MOGEN [30] does not require the conversion of IF to distances and is suitable for large-scale genome structure modeling. GEM [42] considers both Hi-C data and conformational energy derived from knowledge about biophysical models for 3D structure modeling. It used a manifold learning framework, which is aimed at extracting information embedded within a high-dimensional space, in this case the Hi-C data.

Lastly, in the probability-based category, Pastis [25] defined a probabilistic model of IF and casted the 3D inference problem as a maximum likelihood problem. It defined a Poisson model to fit contact data and used an optimization algorithm to solve it. SIMDA3D [46] used a Bayesian approach to infer 3D structures of chromosomes from single cell Hi-C data.

### COMPUTATIONAL MODEL RECONSTRUTION

The GSDB chromosome structure generation was done on three server machines: a x86_64 bit Redhat-Linux server consisting of multi-core Intel(R) Xeon(R) CPU E7-L8867 @ 2.13GHz with 120 GB RAM, x86_64 bit Redhat-Linux server consisting of multi-core Intel(R) Xeon(R) CPU E5649 @ 2.53GHz with 11GB RAM, x86_64 bit Redhat-Linux server consisting of multi-core AMD Opteron(tm) Processor 4284 @ 3.0GHz with 62GB RAM, and a high-performance computing cluster (Lewis) with Linux. Using a high-performance computing (HPC) cluster machine, we allocated 10 cores, 80G of memory, with a time limit of 2 days for each chromosome structure reconstruction task per algorithm. Structures not constructed within 48 hours were terminated.

### DATABASE CONTENT AND USAGE

All the 3D structures in the GSDB have been pre-generated, so that the 3D structure visualization is faster and can be easily downloaded. The steps to navigating the database have been separated into five sections as follows:

i. Browse the database (Figure 1) – Click on “Browse” menu in the navigation bar to load the full list of the Hi-C datasets. Alternatively, users can click on the “Get Started” button on the homepage.
ii. Search the database – The GSDB provides two ways to search for a Hi-C data and its corresponding 3D models:

a. GSDB provides a summary of the information provided in the database through a Summary Pane. By clicking on a property/item in the summary, the user can search the database for all the Hi-C data containing this property and their corresponding 3D structural models. (Figure 2)
b. Users can search the database by typing the keywords about the filename, title of Hi-C data, Hi-C data resolution, project that Hi-C data was generated from (e.g. ENCODE), project ID, and the GEO accession No in the “Search Pane” (Figure 2).
iii. 3D structure visualization and download – To view the details and structures for a Hi-C data, click on the “View” link in the “3D Structure Column” (Figure 3). The data information and visualization tab will be displayed (Figure 4). To show the 3D structure, select the algorithm, dataset, chromosome, and press “Display this Structure” button. The structure will be displayed on the viewer. The modeling parameters and the reconstruction quality (e.g. the Spearman’s correlation between reconstructed distances and expected distances) are reported in the box under the viewer. To compare two structures at the same time, press the “Display Multiple Structures” button. Two structures will be displayed side by side with two distinct options for selecting each visualization’s 3D structure algorithm and dataset (Figure 5). To view a heatmap of the 2D contact matrix used to reconstruct the 3D structure, click the “View Contact Heatmap” button. The heat map can be configured with a variety of helper visualization functions as well as color settings to customize visualization (Figure 6). Users can download the 3D structures by clicking on the “Download” link in the “3D Structure Column” (Figure 3). The normalized Hi-C data used for the 3D structure generation for all the algorithms can also be downloaded by clicking on the “Download” link in the “Normalized Hi-C Data” column (Figure 3).
iv. Evaluation of Structure -- The GSDB contains an evaluation module which permits users to evaluate their own 3D models by comparing model distances to the expected distances of an IF matrix or another 3D model (Figure 7). Upon uploading two PDB files or a PDB file and an IF matrix file and clicking on “Compare” button, users are provided with a collection of evaluation scores including: Spearman Correlation, Pearson Correlation and Root Mean Squared Distance (RMSD).

**Figure 1:**
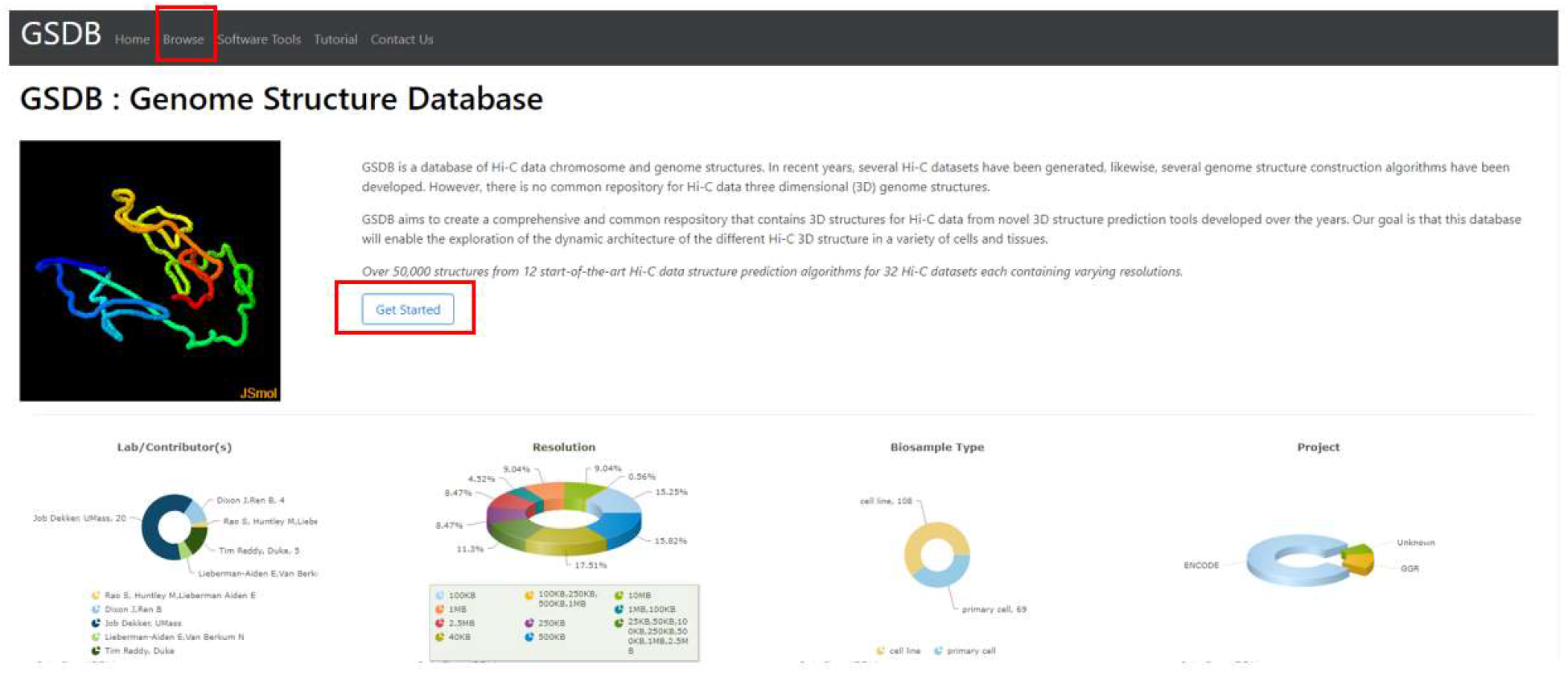
Highlights the two ways to access the database from the homepage. Clicking on the “Browse” menu in the Navigation tab or on the “Get started” button on the home page will load the Database search window.

**Figure 2:**
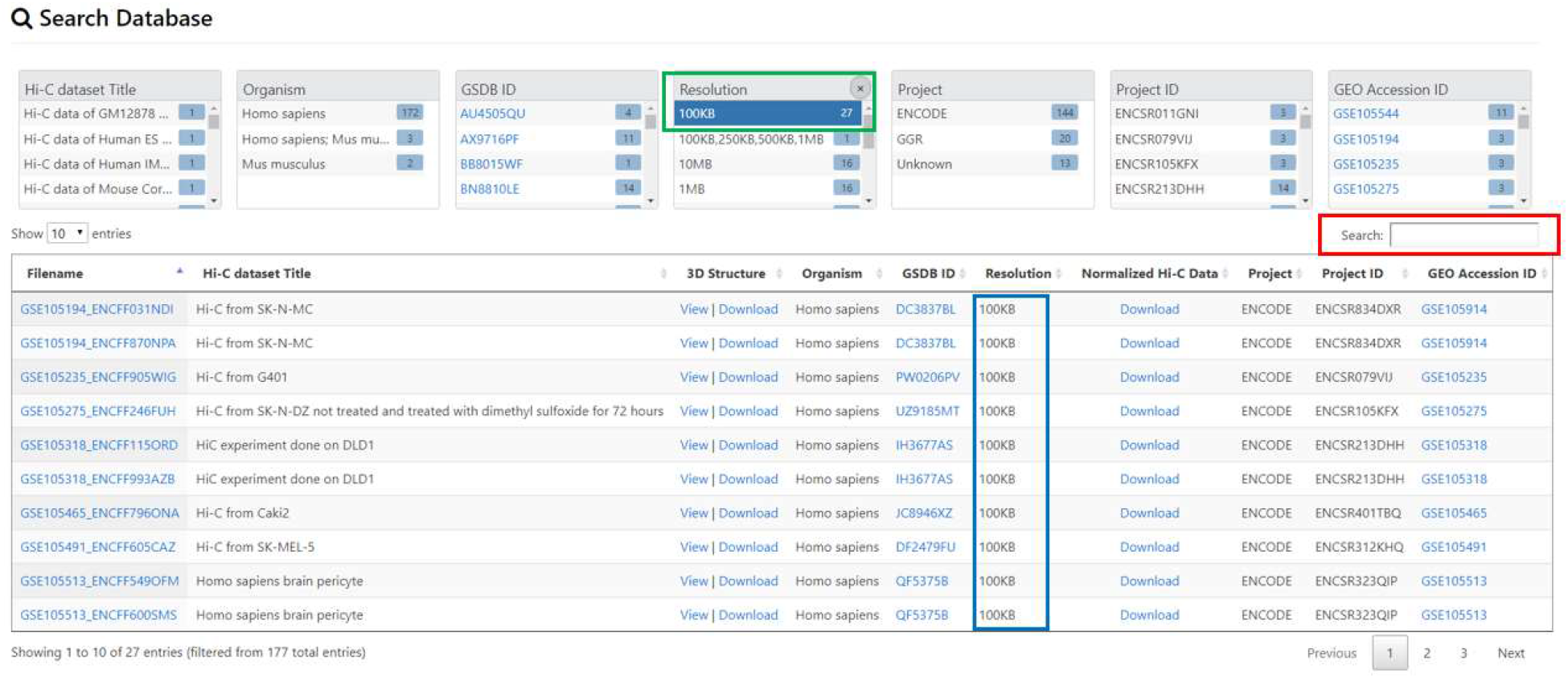
Data search and display. An example of data search using the two approaches for searching. First, search by clicking on an item on the “Summary Pane” highlighted in green. The figure shows when the user clicks on Resolution 100kb, all the datasets with 100kb resolutions are listed. Second, the user can search by typing the key word in the “Search Pane” highlighted in red.

**Figure 3:**
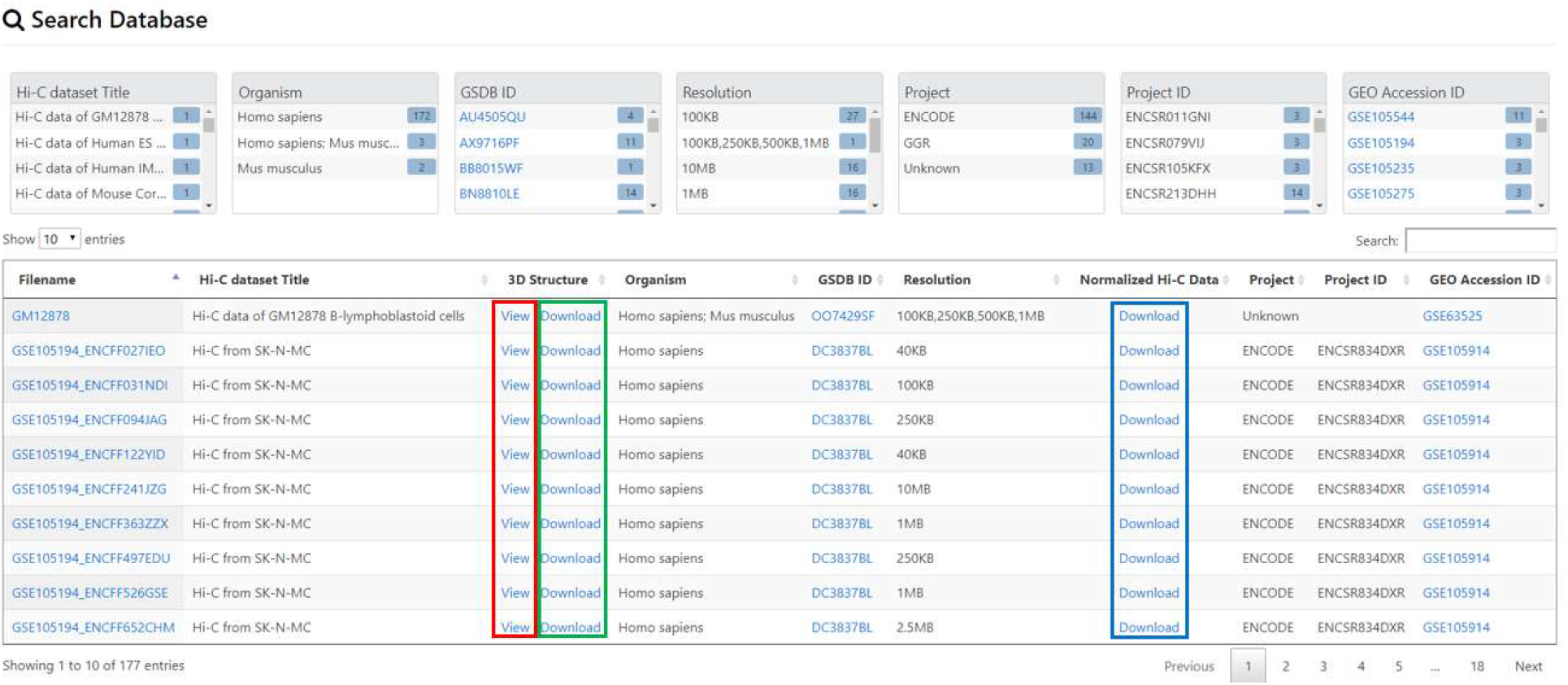
Displaying the database search window. In the “3D Structure” column, highlighted in red is the “View” link to display the 3D structure for a Hi-C data. Highlighted in green is the “Download” link to download the 3D structures constructed by the different algorithms for the Hi-C data. Pressing on the “Download” link will download the 3D structures for all the algorithms for a Hi-C data. In the “Normalized Hi-C Data” column, the “Download” link is highlighted in blue. Pressing on the “Download” link will download the Normalized Hi-C data used for 3D structure construction.

**Figure 4:**
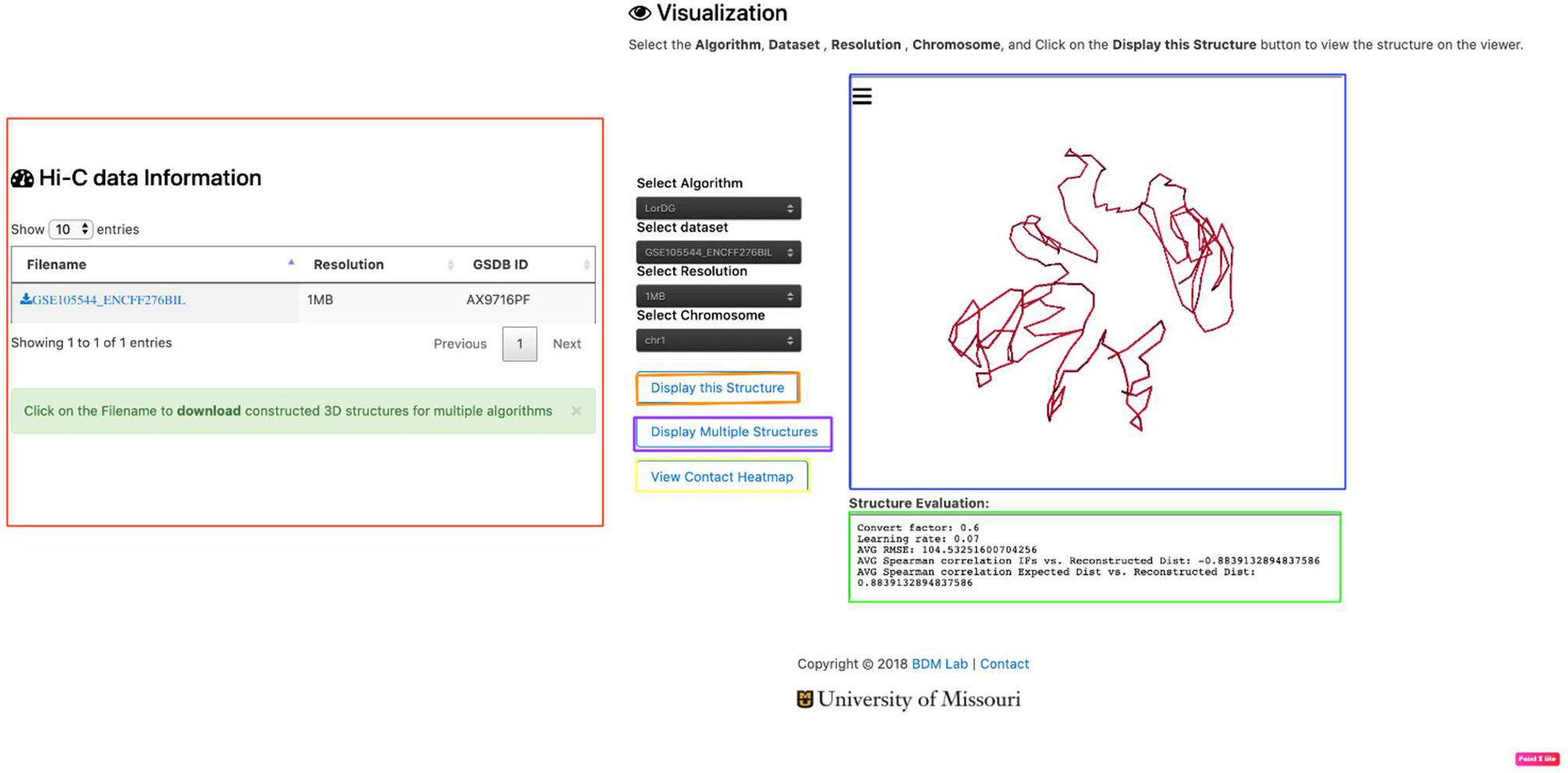
Data visualization. The figure shows the output displayed when a user clicks on the “View link” for the GM12878 dataset. The red highlighted section shows the information about the Resolution(s) available for the Hi-C data. The blue highlighted section displays the structure available for the Hi-C data. The green highlighted section shows the evaluation result available for the Hi-C data. It displays the Spearman Correlation between the output structure and the input Hi-C data, and other evaluation result obtained. To evaluate each 3D structure, we compute the distance Spearman’s correlation coefficient (dSCC) between reconstructed distances and distances obtained from the Hi-C datasets. The value of dSCC is in the range of −1 to +1, where a higher value is better. For distance-based methods, we report the conversion factor (α) used for the IF to distance conversion. For LorDG and 3DMax, which use gradient ascent optimization algorithm, we report the learning rate used for the optimization process. The parameters used by each method to generate 3D structures are available on GSDB GitHub page.

**Figure 5:**
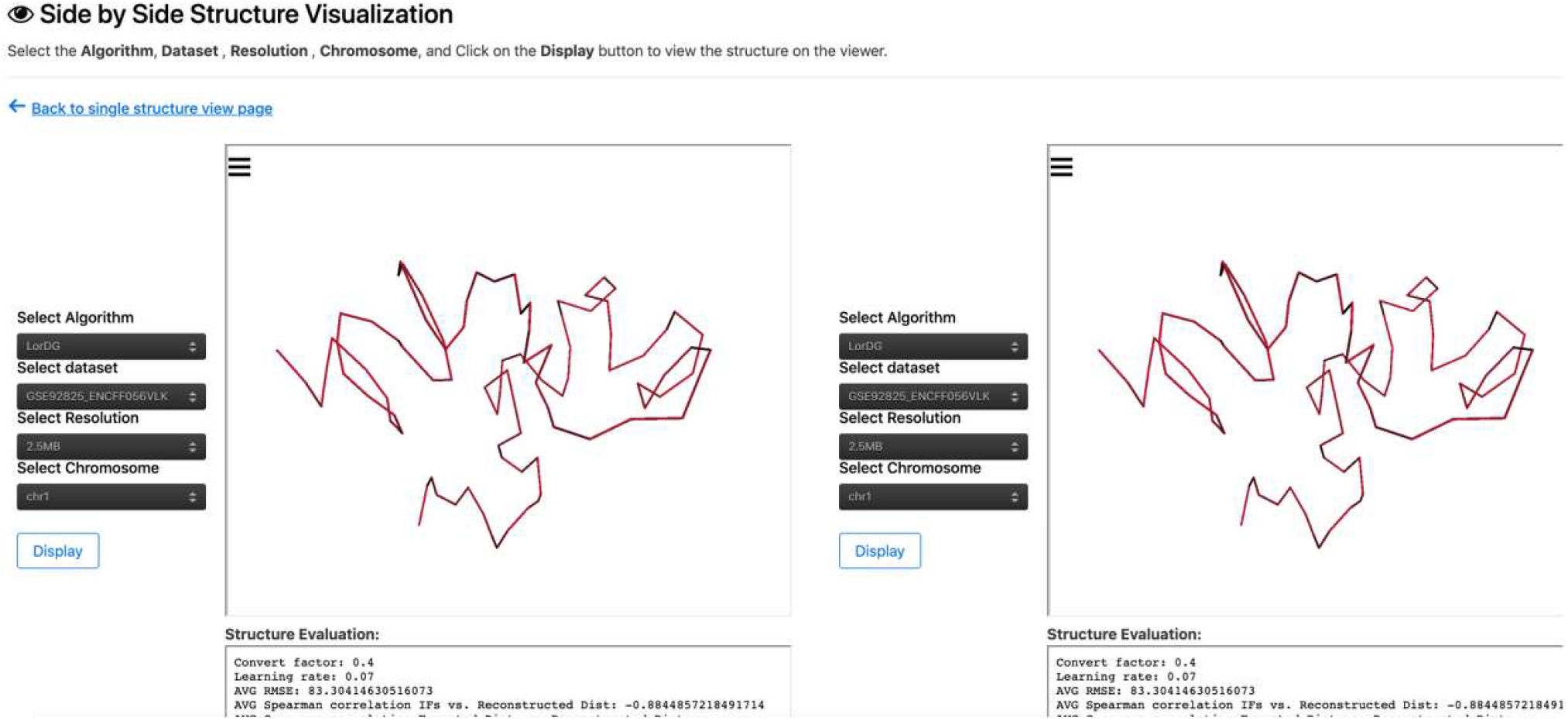
Multiple structure visualization. The figures shows the output displayed when a user clicks the “Display Multiple Structures” button. The multiple structure view permits the comparison of structures using different 3D structure algorithms or different Hi-C contact matrices.

**Figure 6:**
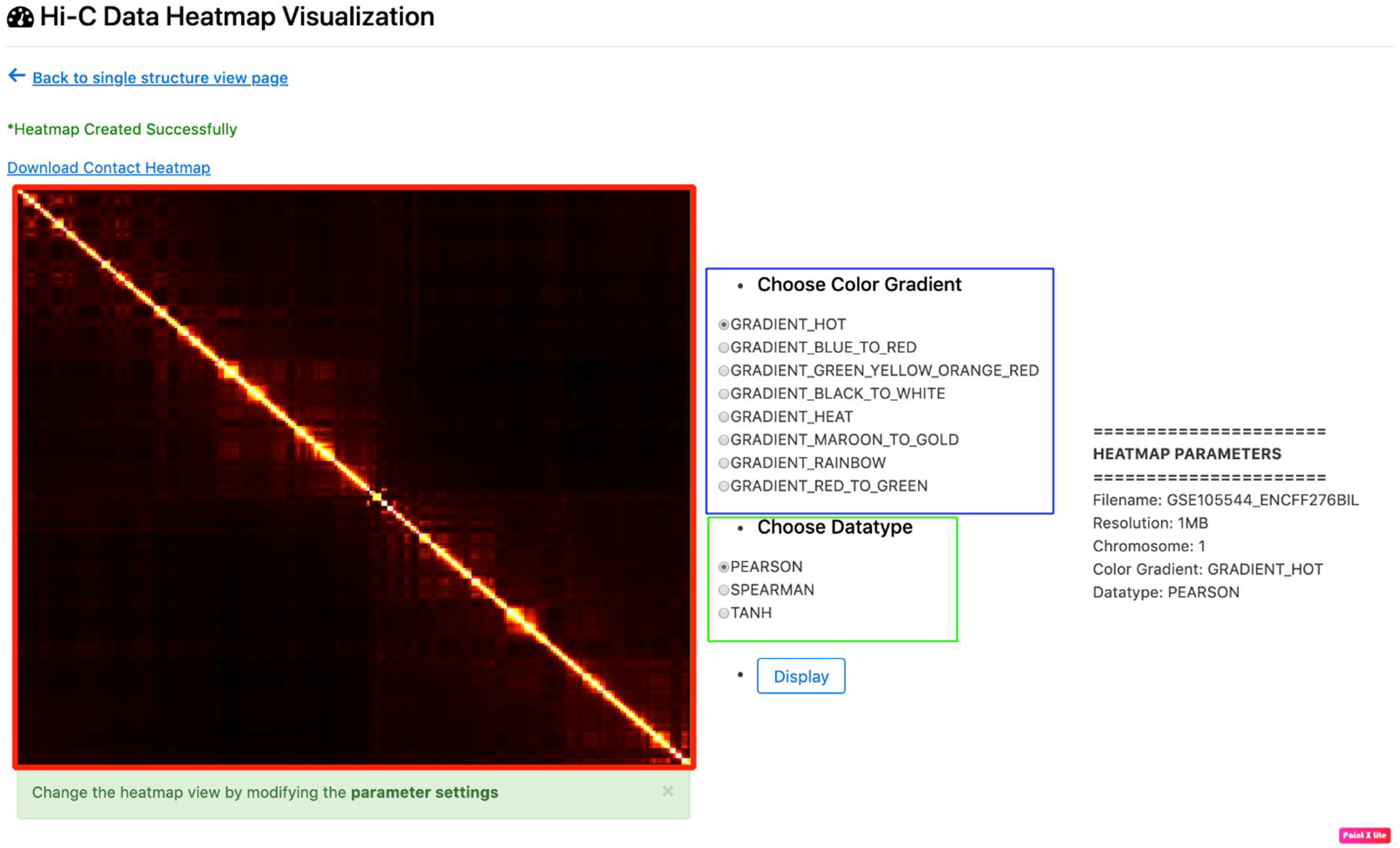
Heat map visualization. The figure shows the output displayed when a user clicks the yellow outlined “View Contact Heatmap” button shown in Figure 4. The figure highlighted in red indicates heat map visualization of the selected 2D chromosomal contact map. The radio buttons outlined in blue display options for the heat map color. The radio buttons outlined in green indicate functions that may be applied to the raw contact matrix prior to heat map construction so as to improve visualization.

**Figure 7:**
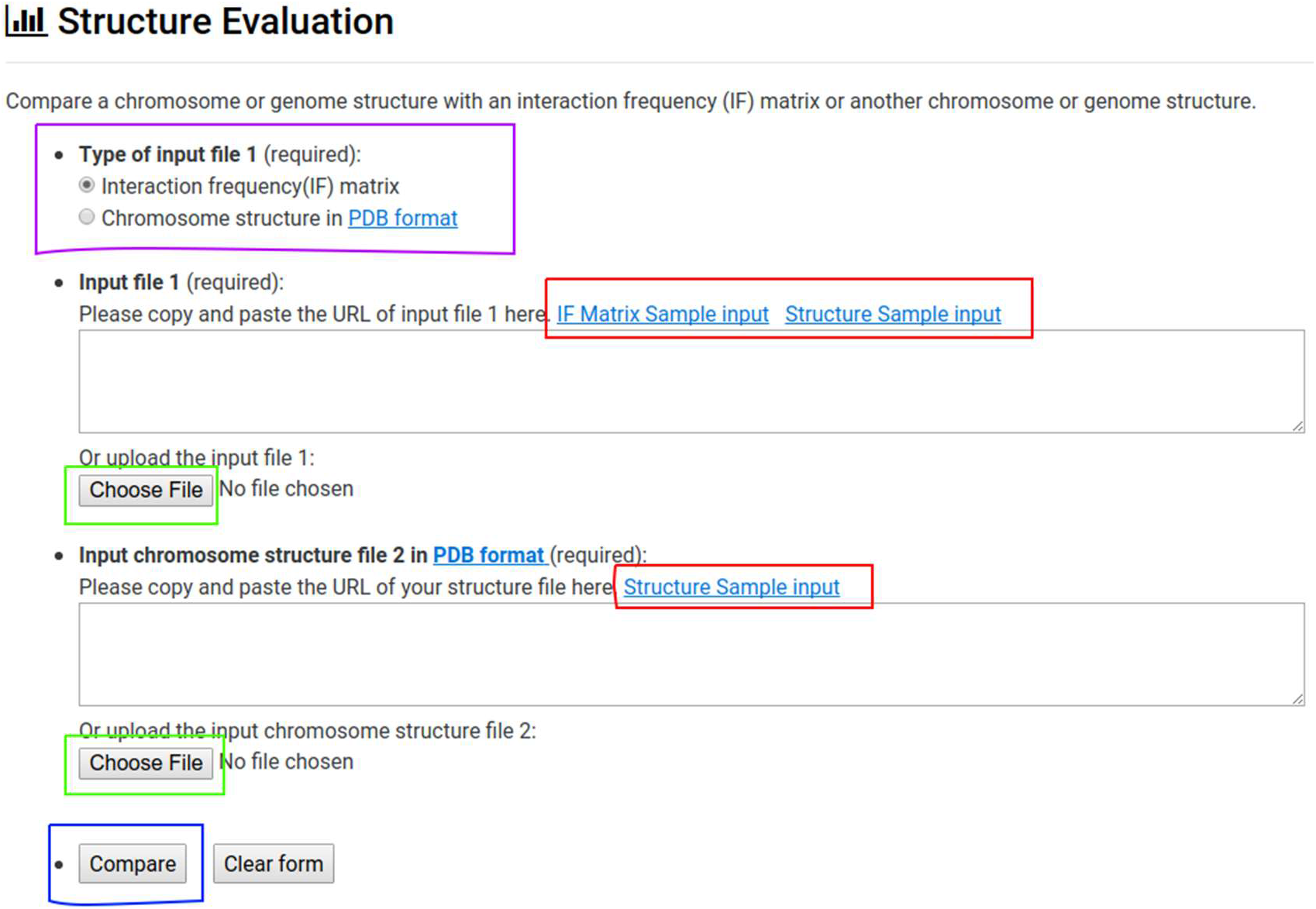
Evaluation. The figure shows the window displayed if a user selects the “Evaluation” tab. The purple box displays the radio buttons which determine whether a comparison will involve 2 structures stored in the Protein Data Bank (PDB) format or a structure in the PDB format and an IF matrix. The green boxes indicate buttons for selecting the files to be compared. The red box denotes links to sample data for testing comparison. The purple box indicates the evaluation button, which will submit the comparison job.

### DISCUSSION AND FUTURE DEVELOPMENT

The GSDB contains 3D structures generated from different Hi-C structure reconstruction algorithms for Hi-C data collected from multiple sources. To the best of our knowledge, it is the first repository for 3D structures generated from multiple Hi-C reconstruction algorithms. Currently, our database contains over 50,000 structures reconstructed for 32 Hi-C datasets by 12 modeling algorithms. The normalized Hi-C dataset used and 3D structures generated from all the algorithms are available to be downloaded. This database will enable the fast and easy exploration of the dynamic architecture of the different Hi-C 3D structure in a variety of cells to improve our understanding of the structural organization of various organisms’ chromosome and genome 3D structures. In addition, we envision that it will be helpful to researchers and scientist to keep track of the performance of the existing approaches for 3D structure construction, and also lead to the development of novel methods that outperform existing approaches. Future directions of the GSDB will include the integration of more algorithms and latest Hi-C datasets generated as the research in 3D structure construction expands.

## AVAILABILITY

GSDB database is freely available at the URL http://sysbio.rnet.missouri.edu/3dgenome/GSDB. Scripts and the parameters used for the 3D structure generation for each algorithm are available at https://github.com/BDM-Lab/GSDB

## ACKNOWLEDGEMENTS

The computation for this work was performed on the high-performance computing infrastructure provided by Research Computing Support Services and in part by the National Science Foundation under grant number CNS-1429294 at the University of Missouri, Columbia.

